# Mtg16-dependent repression of E protein activity is required for early lymphopoiesis

**DOI:** 10.1101/2020.07.24.220525

**Authors:** Pankaj Acharya, Shilpa Sampathi, David K. Flaherty, Brittany K. Matlock, Christopher S. Williams, Scott W. Hiebert, Kristy R. Stengel

**Affiliations:** Department of Biochemistry, Vanderbilt University School of Medicine, Nashville, Tennessee 37232, USA; Vanderbilt Flow Cytometry Shared Resource; Program in Cancer Biology, Vanderbilt University, Nashville, TN, USA; Department of Medicine, Division of Gastroenterology, Vanderbilt University Medical Center, Nashville, Tennessee 37232, USA; Vanderbilt-Ingram Cancer Center, Vanderbilt University School of Medicine, Nashville, Tennessee 37027, USA

## Abstract

The ETO/MTG family of transcriptional co-repressors play a key role in adult stem cell functions in various tissues. These factors are commonly found in complex with E proteins such as E2A, HEB, and Lyl1 as well as PRDM14 and BTB/POZ domain factors. Structural studies identified a region in the first domain of MTGs that is conserved in the *Drosophila* homologue Nervy (Nervy Homology Domain-1, or NHR1) that is essential for ETO/MTG8 to inhibit E protein-dependent transcription. The Cancer Genome Atlas (TCGA) identified cancer associated single nucleotide variants (SNVs) near the MTG16:E protein contact site. We tested these SNVs using sensitive yeast two-hybrid association assays, which suggested that only P209T significantly affected E protein binding. We then used CRISPR-Cas9 and homology directed DNA repair to insert P209T and a known inactivating mutation, F210A, into NHR1 of *Mtg16* in the germ line of mice. These mice developed normally, but in competitive bone marrow transplantation assays, the F210A-containing stem cells failed to contribute to lymphopoiesis, while P209T mutant cells were reduced in mature T cell populations. High content fluorescent activated analytical flow cytometry assays identified a defect in the multi-potent progenitor to common lymphoid progenitor transition during lymphopoiesis. These data indicate that the cancer associated changes are likely benign polymorphisms, and the MTG:E protein association is required for lymphopoiesis, but less important for myelopoiesis and stem cell functions.

## Introduction

The myeloid translocation gene (MTG) family of transcriptional corepressors are frequent targets of chromosomal translocation in myeloid leukemias and have emerged as critical regulators of cell fate decisions in hematopoiesis, neurogenesis, and the intestines (Chyla et al., 2008; Fischer et al., 2012; Koyano-Nakagawa and Kintner, 2005; Parang et al., 2015; Peterson and Zhang, 2004; Steinauer et al., 2017). MTG proteins generally serve as scaffolds to recruit HDAC-containing corepressor complexes to a variety of sequence-specific transcription factors, the most well characterized of which are the class I bHLH transcription factors known as E proteins (de Pooter and Kee, 2010). The MTG family consists of three proteins: MTG8 (ETO, RUNX1T1, CBFA2T1), MTG16 (CBFA2T3, ETO2), and MTGR1 (CBFA2T2), each containing four homology regions (NHR1-4) with high conservation to the *Drosophila* protein, Nervy (Erickson et al., 1992; Gamou et al., 1998; Kitabayashi et al., 1998; Miyoshi et al., 1993).

The first conserved domain, nervy homology region 1 (NHR1), is also homologous to hTAF130 and hTAF105, which are components of the TFIID general transcription factor complex (Erickson et al., 1994). It is this homology region that mediates most contacts with sequence-specific transcription factors including E proteins (Moore et al., 2008; Park et al., 2009; Zhang et al., 2004), PRDM14 (Nady et al., 2015), GFI-1/1B (McGhee et al., 2003), and ZBTB4/38/33 (Barrett et al., 2012). The second homology region, NHR2, forms anti-parallel heterotetramers (Liu et al., 2006) potentially containing all three MTG family members. This highly stable tetramer formation brings four NHR3/4 domains into close proximity at bound loci. While NHR3 is the least conserved domain and does not have a well-defined function, it is NHR4 that is responsible for recruitment of NCOR1/SMRT/HDAC3 corepressor complexes as well as other chromatin regulators to facilitate gene repression (Amann et al., 2001; Gelmetti et al., 1998; Lutterbach et al., 1998a; Lutterbach et al., 1998b; Wang et al., 1998).

While mouse models revealed critical roles for all three Mtg family members in stem cell function and cellular differentiation in a variety of tissue types (Amann et al., 2005; Baulies et al., 2020; Calabi et al., 2001; Nady et al., 2015; Williams et al., 2013), only Mtg16 has been implicated in the regulation of normal hematopoietic stem cell (HSC) function and hematopoietic development (Chyla et al., 2008; Fischer et al., 2012; Hunt et al., 2011). Although *Mtg16*^*−/−*^ mice were viable, *Mtg16*^*−/−*^ HSPCs exhibited defects in proliferation and self-renewal capacity, which sensitized *Mtg16*^*−/−*^ animals to erythropoietic stress (Chyla et al., 2008) and prevented transplantation of *Mtg16*-deficient bone marrow (Fischer et al., 2012). *Mtg16* deletion also impaired T cell development both *in vivo* and *in vitro*, and while re-expression of wild-type Mtg16 could complement that defect, an NHR1 domain deletion mutant could not (Hunt et al., 2011).

E proteins are the best-characterized interacting proteins with the NHR1 domain of MTG family members. There are four ubiquitous class I bHLH proteins (a.k.a. E proteins): HEB/TCF12, E2-2/TCF4, and two E2A proteins (E47 and E12), which form homo- or heterodimers to bind E box sequences (CANNTG) and activate gene expression (de Pooter and Kee, 2010). E proteins can also dimerize with the more tissue-specific class II bHLH proteins including the hematopoietic transcription factors, SCL/TAL1 and LYL1, which are both targeted by translocations in T-cell acute lymphoblastic leukemia (Souroullas et al., 2009). Proteomic studies have identified MTG proteins in complex with SCL/TAL1 and LYL1 in addition to E proteins (Sun et al., 2013), and MTG16 was shown to regulate the activity of SCL/TAL1:E2A complexes in hematopoietic progenitor cell populations (Cai et al., 2009; Schuh et al., 2005; Stadhouders et al., 2015).

Here, we screened cancer-associated point mutations located within the NHR1 domain of Mtg16 for the ability to disrupt E protein binding and Mtg16-mediated transcriptional repression. While none of these mutations impaired Mtg16 repressor function to the same extent as the previously identified Mtg16^F210A^ mutation, we did identify an additional mutant, Mtg16^P209T^, which exhibited a moderate impairment of Mtg16 repressor activity in reporter assays. To further examine the impact of these mutations on Mtg16 function, we used CRISPR-based genome editing to introduce the F210A and P209T mutations into the NHR1 domain of the murine Mtg16 locus. As the loss of Mtg16 results in well-characterized hematopoietic defects, we utilized competitive bone marrow transplantation assays to assess the impact of these Mtg16 mutations on overall Mtg16 function. While significant defects in hematopoiesis were observed in Mtg16^F210A^ marrow, they were not as severe as those observed with *Mtg16* deletion, suggesting that Mtg16 likely represses transcription factors in addition to E proteins to fully regulate hematopoiesis. Moreover, the most pronounced effects of Mtg16^F210A^ were on lymphoid rather than myeloid cell populations, suggesting that the ability of Mtg16 to regulate E protein targets is particularly important for lymphopoiesis. Finally, the moderate effect of the P209T mutation on the ability of Mtg16 to repress E protein activity did not result in substantial changes in hematopoietic cell development. The one exception to this was a decrease in the most mature T cell populations, underscoring that the ability of Mtg16 to regulate E protein-mediated transcription is especially important for lymphoid development.

## Materials and Methods

### Mice

Animals were housed under pathogen-free conditions. All experiments were conducted in accordance with an Institutional Animal Care and Use Committee-approved protocol under guidelines set forth by Vanderbilt University Medical Center. Mtg16 mice were generated previously. *Mtg16*^*F210A*^ and *Mtg16*^*P209T*^ mice were generated by cytoplasmic injection of 50 ng/μl gRNA, 200 ng/μl of 119 bp ssDNA donor oligonucleotide, and 10 ng/μl *Cas9* mRNA. 25 μl was injected per mouse embryo, embryos transferred to recipient mice and tails from resultant pups screened for successful editing by PCR. gRNA sequence: TTTGTTATCCCTTTTCTGAAGG. F210A donor oligo sequence: TTCATGCCAAGCTCCAGGAAGCCACCAACTTTCCACTGAGGCCGTTTGTTATCCCTG CTCTGAAAGTAAGCGAAAGCAGCACCTTCCAGGCAACAGGGATGGTGTACACAAAGGCCCTG. P209T donor oligo sequence: GTTTCATGCCAAGCTCCAGGAAGCCACCAACTTTCCACTGAGGCCGTTTGTTATAACTTTTCTGAAAGTAAGCGAAAGCAGCACCTTCCAGGCAACAGGGATGGTGTACACAA AGGCCC.

### Genotyping

*Mtg16F210A and Mtg16P209T* were each genotyped using mutation-specific reverse primer. The first reverse primer recognized the wild-type Mtg16 sequence and the second specifically recognized the mutant allele. Primers: F210A/P209T-F: AGCACCTGCCCCCGGCGTGCG, F210A-wt-R:TTGCCTGGAAGGTGCTGCTTTCGCTTACCTTCAGAAA, F210A-mut-R:TTGCCTGGAAGGTGCTGCTTTCGCTTACTTTCAGAGC, P209T-wt-R: TTGCCTGGAAGGTGCTGCTTTCGCTTACCTTCAGAAAAGGG, P209T-mut-R: TTGCCTGGAAGGTGCTGCTTTCGCTTACTTTCAGAAAAGTT.

### Bone Marrow Transplants

Bone marrow was isolated from 8-week-old wild-type, *Mtg16*^*−/−*^, *Mtg16*^*F210A*^ and *Mtg16*^*P209T*^ animals as well as wild-type CD45.1 competitors (NCI B6-Ly5.1/Cr). 1 million CD45.2^+^ donor cells were mixed with 1 million CD45.1^+^ competitor cells and the 50:50 mix was transplanted by tail vein injection into lethally irradiated B6-Ly5.1 recipient mice. 12 weeks post-transplant, animals were euthanized and bone marrow, spleen, and thymus were recovered in order to assess hematopoietic cell populations.

### Spectral Flow Cytometry

Bone marrow was isolated by flushing and red cell lysis performed by resuspending pellet in 1 ml of Buffer EL (Qiagen) and incubating on ice for 15 minutes. Cells were then resuspended at a concentration of 5 million cells/500 μl in PBS + 0.5% BSA. Bone marrow cells were stained with a panel of 20 antibodies (see Table S1) for 1 hour at 4°C. For lymphoid tissues (thymus and spleen) single cell suspensions were obtained by pressing the tissue through a 70-μM strainer. Following erythrocyte lysis, cells were resuspended at 2 million cells/200 μl in PBS + 0.5% BSA and stained with an 18-antibody lymphoid panel (see Table S1) for 1 hour at 4°C. Post-staining, cells were washed with 1 ml PBS + 0.5% BSA, resuspended in 500 μl and acquired on a Cytek Aurora spectral flow cytometer. Data analysis was performed using FlowJo software (version 10.7.0, BD).

### Plasmids

Generation of pCMV5-Myc-Mtg16 point mutants (F210A and R220A) was described previously (Hunt et al., 2011). Other cancer associated Mtg16 point mutants (http://www.cbioportal.org/public-portal/) were generated by site-directed mutagenesis using iProof DNA polymerase (Bio-Rad). The oligonucleotides used for mutagenesis are listed in Table S2.

### Luciferase Reporter Assays

Luciferase assays were performed as previously described (Hunt, 2011). Briefly, 2 × 10^5^ 293T cells were seeded per well in 6-well dishes. After 24 hours the cells were cotransfected with 100 ng of MTG16 (wild type or mutants), E47, renilla luciferase and E-box reporter plasmids using polyethylenimine (PEI). After 48 hours the cells were lysed with passive lysis buffer (Dual-Luciferase reporter assay system; Promega) and the luciferase activity was measured on a BD Pharmingen Monolight 3010 luminometer. The transfections were performed in triplicates, and the values were normalized to that of renilla luciferase internal control.

### Yeast-2-Hybrid Assays

*S. cerevisiae* (L40, H83) were transformed with HEB and vector or MTG16 constructs. Three independent single colonies were cultured overnight. The next day those cells were inoculated into fresh medium and incubated for further 4-5 hours. Then 5 × 10^6^ cells were used to perform β galactosidase activity using Tropix Gal-Screen System (Applied Biosystems). The readings were normalized to MTG16 wild type activity. Three independent colonies serve as triplicates.

### Co-immunoprecipitation

4 × 10^5^ 293T cells were seeded per well in 6-well dishes. After 24 hours the cells were transfected with 200 ng of MTG16 (wild type or mutants) using polyethylenimine (PEI). For E47-MTG16 IP, 200 ng of E47 was transfected as well. After 48 hours, the cells were harvested in buffer containing 20 mM Tris (pH 7.4), 100 mM NaCl, 1% Triton X-100, 1 mM EDTA and protease inhibitors. The lysates were sonicated and cleared by centrifugation. Anti E47 (sc-763; Santa Cruz) and Anti HEB (sc-357; Santa Cruz) antibodies were used for E47-MTG16 and HEB-MTG16 coimmunoprecipitations, respectively. The complexes were collected using protein A/G agarose beads (Santa Cruz). The coimmunoprecipitating proteins were detected by western blot using anti-c-Myc 9E10 antibodies.

## Results

### NHR1-associated point mutations impair Mtg16-E protein binding and Mtg16-mediated repression

The Cancer Genome Atlas (TCGA) has reported numerous single nucleotide variants across the NHR1 domain of MTG family members; however, whether these genomic changes are true mutations affecting MTG function remains to be explored. Therefore, we screened cancer-associated variants for their ability to disrupt binding of Mtg16 to the E protein, HEB, by yeast-two-hybrid (Y2H) assay (Fig. 1A). Deletion of the entire NHR1 domain resulted in complete disruption of Mtg16-HEB binding, and the previously characterized point mutant, F210A, resulted in a comparable loss of the Mtg16-HEB association. In contrast, most cancer-associated variants had little effect on E protein binding. One notable exception was Mtg16^P209T^, which was significantly impaired in its ability to bind HEB. We then used reporter assays to determine the ability of Mtg16 variants to repress E protein-mediated transcription (Fig. 1B). While wild-type Mtg16 efficiently repressed E protein-mediated transactivation, the F210A point mutant was unable to repress E2A-mediated transcription. Consistent with Y2H assays, most cancer-associated variants remained fully functional, though two mutants (P209T and P205A) exhibited a partial loss of repressor activity. Therefore, we focused on these mutations for subsequent functional analyses.

**Figure 1.**
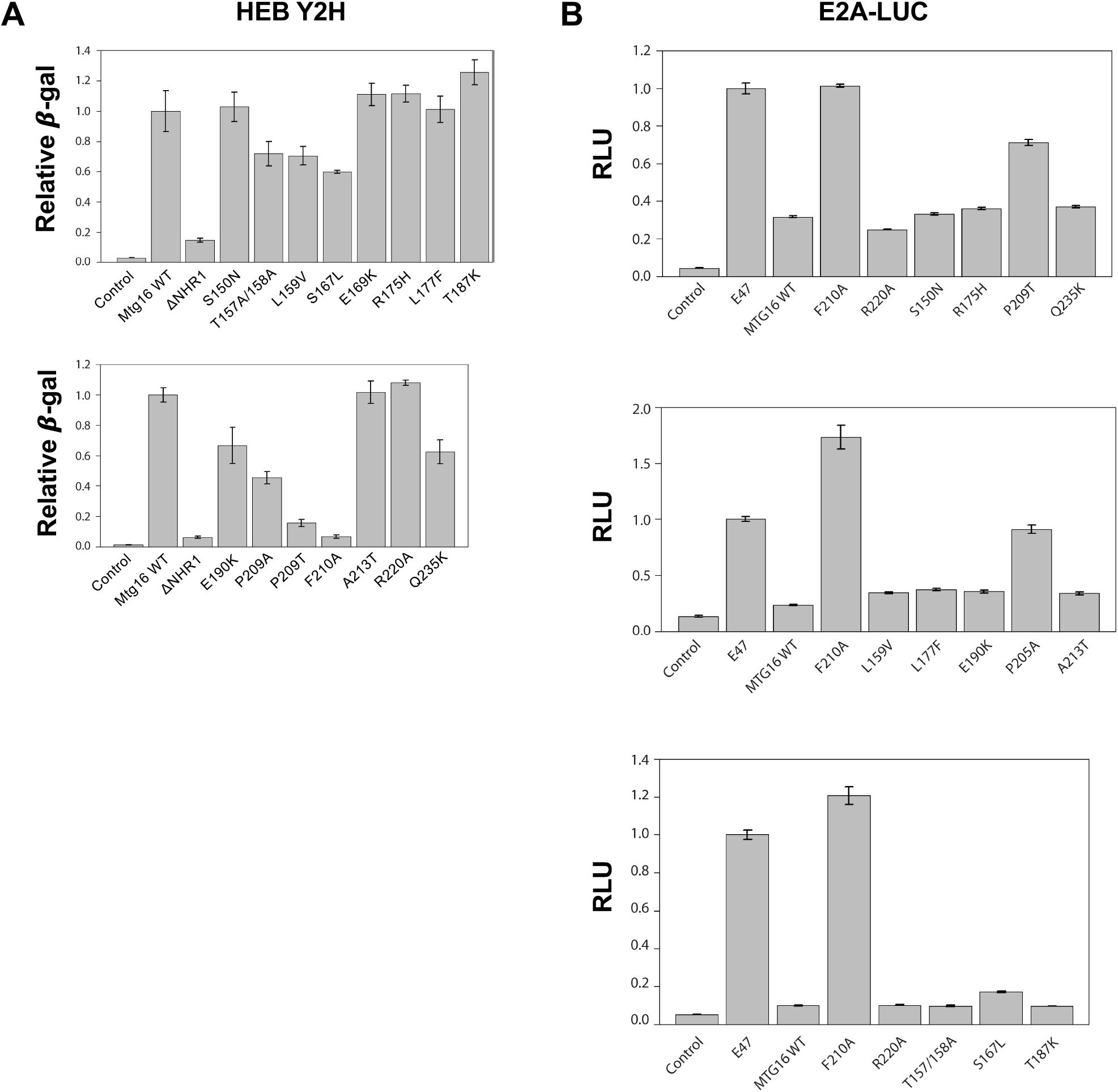
Screening of cancer-associated NHR1 point mutations identified effects on E protein binding and repression. (A) The indicated point mutations in the NHR1 domain of Mtg16 were screened for their ability to impair HEB binding by Y2H assay. Yeast-two-hybrid assays were performed on yeast transformed with the indicated Mtg16 point mutant fused to LexA DNA-binding domain and a fragment of HEB (aa9-642) fused to the Gal4 activation domain. Binding dependent β-galactosidase expression was quantified using the Tropix Gal-Screen System and β-gal activity was normalized to that produced by wild-type Mtg16. Three independent transformants are shown. (B) The indicated point mutations in the NHR1 domain of Mtg16 were screened for their ability to impair E2A-mediated reporter gene repression. 293T cells that were transfected with Myc-Mtg16 were also transfected with E47, the E-box-Luc reporter plasmid, and renilla luciferase control construct. Relative firefly luciferase activity was quantified, normalized to renilla activity and plotted relative to vector control samples.

We used Y2H assays to determine the ability of Mtg16 mutants to bind to a second E protein, TCF4, and observed that similar to HEB binding, Mtg16^F210A^ was unable to bind TCF4, while Mtg16^P209T^ and Mtg16^P205A^ showed partial disruption of TCF4 binding (Fig. 2A). In contrast, when we asked whether these mutations would disrupt binding to Zbtb38, a distinct transcription factor reported to bind NHR1, we found that neither the F210A nor the P209T mutations affected the ability of Mtg16 to bind Zbtb38 (Fig. 2B), suggesting that these mutations selectively disrupted Mtg16-E protein interactions. As Y2H assays were performed with peptide fragments of HEB and TCF4, we next performed co-immunoprecipitation assays following overexpression of Mtg16 mutants in 293T cells to determine the effect of point mutations on the association of Mtg16 and full-length E47 or HEB in mammalian cells. While both F210A and P209T mutations appeared to completely disrupt the association of Mtg16 with E47 (Fig. 2C), some residual binding to HEB was observed (Fig. 2D), which is consistent with Mtg proteins making multiple contacts with E proteins, including some contacts mediated by NHR2 (Sun et al., 2013). Alternatively, this could reflect co-purification facilitated by endogenous Mtg proteins that are associated with mutants through tetramer formation.

**Figure 2.**
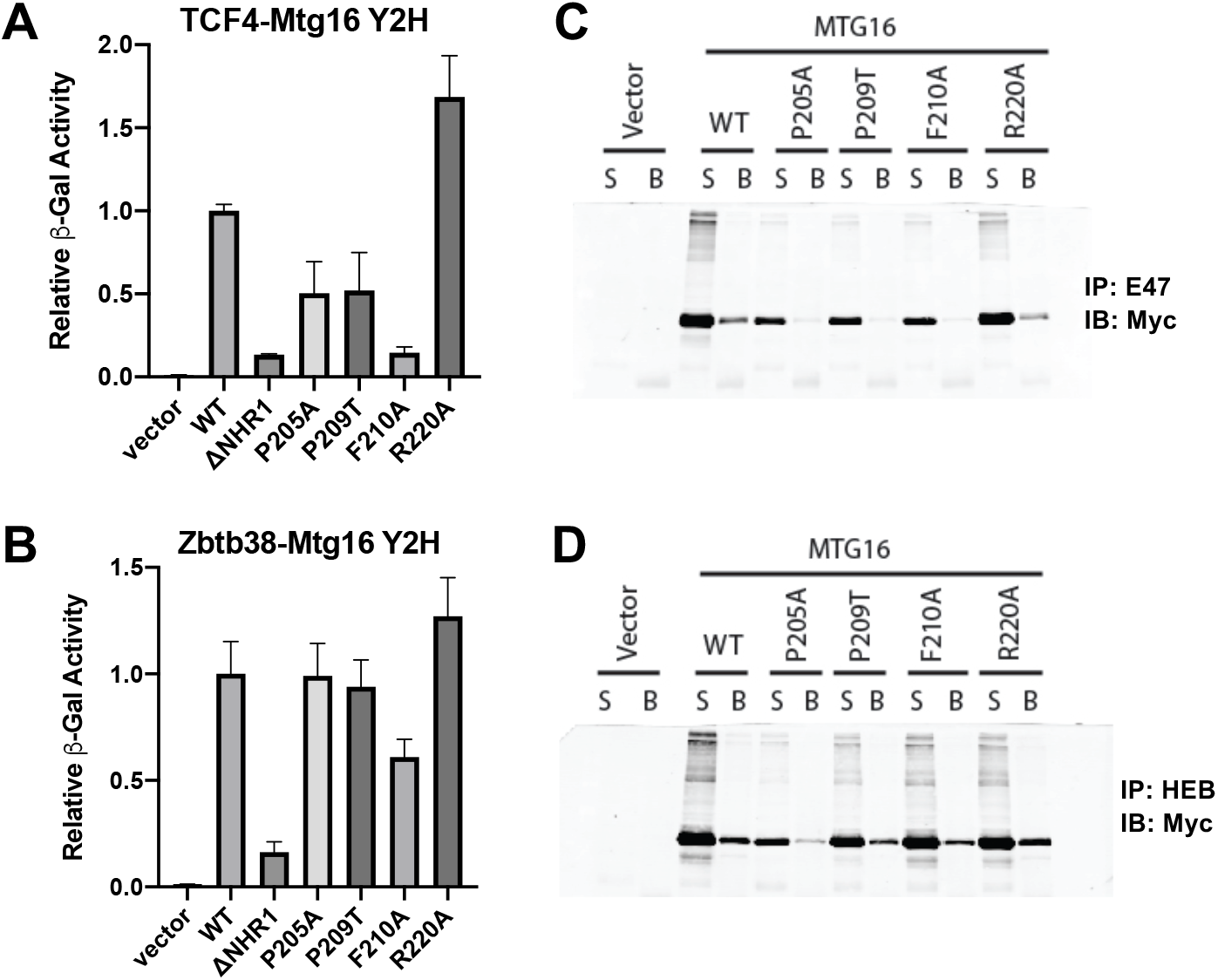
Point mutations in NHR1 disrupt E protein binding and repression. Y2H assays were performed as in Figure 1, using TCF4(aa1-652)-Gal4-AD (A) or Zbtb38-Gal4-AD (B) as prey. (C) Whole cell lysates from 293T cells transfected with the indicated Myc-Mtg16 mutant were immunoprecipitated using an antibody to the E protein, E47. Protein association was detected by immunoblotting with anti-Myc. S=supernatant B=beads (D) Lysates from C were also immunoprecipitated with anti-HEB and protein association detected by anti-Myc.

### Failure of Mtg16 to regulate E proteins results in a loss of splenic lymphoid populations

In order to determine how complete (F210A) or partial (P209T) disruption of E protein repression by Mtg16 contributes to overall Mtg16 function, we used CRISPR-based genome editing to insert the F210A and P209T point mutations into the endogenous *Mtg16* locus in mice. Deletion of *Mtg16* had profound effects on hematopoietic stem cell function and blood cell development (Chyla et al., 2008; Fischer et al., 2012; Hunt et al., 2011). Therefore, we performed competitive bone marrow transplants using 50% wild-type CD45.1 bone marrow and 50% CD45.2 marrow from either wild-type negative control animals (n=8), *Mtg16*-deleted positive control animals (n=7), Mtg16^F210A^ animals (n=8), or Mtg16^P209T^ animals (n=8). Twelve weeks following transplant, hematopoietic tissues were harvested and the relative contribution of CD45.2^+^ marrow to numerous hematopoietic cell populations was determined by high content spectral flow cytometry. Using a spectral cytometer allowed us to stain cells with antibody panels targeting up to 20 cell surface proteins, thus providing a more comprehensive definition of hematopoietic cell populations than can be accomplished by traditional flow cytometry approaches that are limited due to the conventional methods used to account for the spectral overlap of fluorophores.

We first asked whether Mtg16 mutant proteins had any alterations in mature splenocyte populations. For splenocyte spectral flow analysis, down-sampling was performed and data from all mice merged prior to t-SNE dimensionality reduction on the basis of unmixed lineage parameters (B220, CD3, Gr1, CD11b, Ter119) using FlowJo software. Manually gated B220+ B cell, CD3+ T cell, Ter119+ erythroid, and CD11b+ myeloid populations were identified within the generated t-SNE plot (Fig. 3A). We then deconvolved merged data based on genotype and displayed CD45.1^+^ vs. CD45.2^+^ cell populations on this same t-SNE plot for each Mtg16 mutant (Fig. 3B). Wild-type competitor cells (CD45.1^+^) are displayed in red, while CD45.2+ donor cells are displayed in blue, such that any population showing an overrepresentation of red events suggests that donor cells were outcompeted by wild-type control cells.

**Figure 3.**
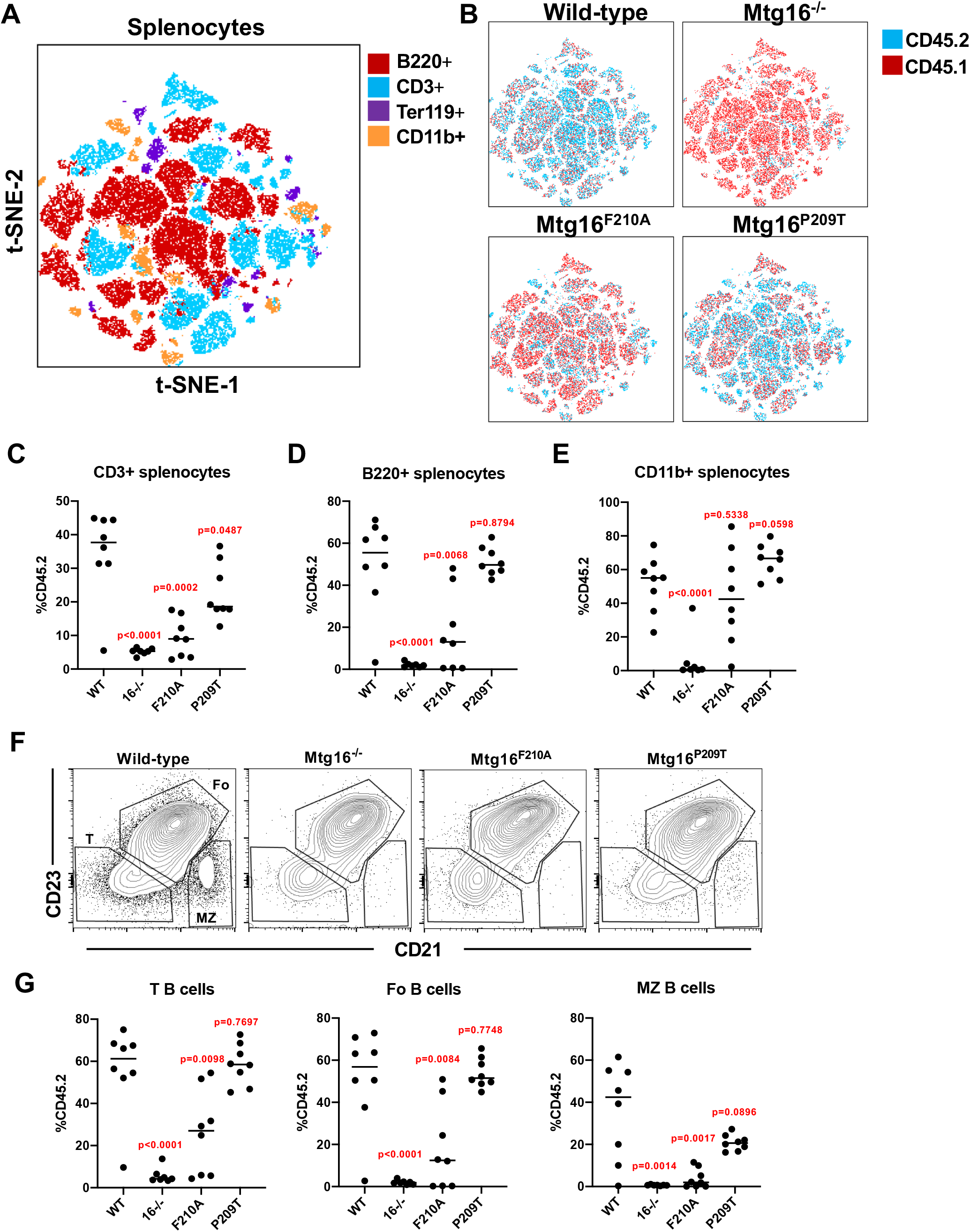
Mtg16 point mutations alter lymphoid populations in the spleen. (A) 12 weeks post-transplant with 50% wild-type CD45.1 marrow and 50% CD45.2 marrow of the indicated genotype, spleens were harvested and spectral flow cytometry performed to identify mature cell populations in the spleen. T-SNE dimensionality reduction was performed on a concatenated file containing 3000 downsampled events from each mouse across all genotypes. Manually gated B220^+^, CD3^+^, Ter119^+^, and CD11b^+^ populations projected onto the resulting plot. (B) Each genotype was gated from the concatenated file and the relative contribution of CD45.1^+^ and CD45.2^+^ cells were displayed on the t-SNE plot. (C-E) The percentage of CD45.2^+^ CD3^+^ (C), B220^+^ (D), and CD11b^+^ (E) splenocytes were quantified for each mouse. (F) CD45.2^+^B220^+^ splenic B cells were further gated based on CD21 and CD23 expression to classify cells as transitional (T), follicular (Fo), or marginal zone (MZ) B cells. Quantification of these populations are presented (G).

For wild-type transplanted animals, all splenic populations showed a relatively equal distribution of CD45.1^+^ and CD45.2^+^ cells (Fig. 3B, upper left). Consistent with previously published studies, animals transplanted with *Mtg16*^*−/−*^ marrow showed a dramatic overrepresentation of CD45.1^+^ cells in all splenic populations suggesting that *Mtg16*-deficient myeloid, B- and T-cells were all rapidly outcompeted. In contrast, *Mtg16*^*F210A*^ transplanted animals showed a dramatic over-representation of CD45.1^+^ cells in B- and T- cell populations, while CD11b+ myeloid populations still had a substantial contribution from CD45.2^+^ mutant cells (see patches of blue that remain, Fig. 3B, lower left). Finally, the spleens of *Mtg16*^*P209T*^ transplanted animals showed a specific selection for CD45.1 wild-type cells only in CD3+ T-cell populations\], suggesting that complete repression of E protein-mediated transcription may be particularly important for T cells. We further quantified the contribution of CD45.2^+^ marrow to CD3^+^, B220^+^, and CD11b+ splenic populations by manual gating, and found the same trends whereby Mtg16 loss dramatically impacted all cell populations, Mtg16^F210A^ resulted in a reduction of B- and T- cell populations, and Mtg16^P209T^ had a specific effect on CD3^+^ T-cell populations (Fig. 3C-D).

We further analyzed splenic B cell populations based on CD21 and CD23 expression to define transitional (T), follicular (Fo), and marginal zone (MZ) B cell populations (Fig. 3F). Both *Mtg16*^*−/−*^ and *Mtg16*^*F210A*^ transplanted animals exhibited a significant reduction in the percentage of CD45.2^+^ T-, Fo-, and MZ-B cells. In contrast, *Mtg16*^*P209T*^ transplanted animals showed no reduction in CD45.2^+^ cell contribution to T- or Fo-B cell populations, and though CD45.2^+^ MZ-B cells were trending downward, it did not reach significance (Fig. 3G).

### The effect of Mtg16^F210A^ on B cell development occurred prior to B-cell lineage commitment in the marrow

In order to determine how the loss of Mtg16 and the presence of the F210A mutation caused a reduction in splenic B cell populations, we examined B cell development within the bone marrow. B-cell development is tightly linked to recombination of the immunoglobulin locus to create a functional B cell receptor (BCR). Recombination of the heavy chain locus occurs first and coincides with a transition from a CD43^+^ B cell to a CD43^−^ B cell. CD43 positive and negative populations can be further divided into “Hardy Fractions”, which more specifically define B cell progenitor populations (Hardy et al., 1991).

We first limited our analysis to B cell populations within the marrow by excluding Ter119^+^, CD3^+^, Gr1^+^ and CD11b^+^ populations and gating specifically on B220^+^ cells. We then performed t-SNE dimensionality reduction as described above, on the basis of unmixed B cell markers (CD24, BP-1, IgD, IgM, IL7R, and CD43; Figure 4A, upper left). Manually gated Hardy fractions (Fig. S1) were projected onto the t-SNE plot and color-coded to reflect Hardy Fractions A-F (Fig. 4A, lower left). Heatmaps depict the relative expression of the cell surface markers used to define Hardy Fractions (Fig. 4A, right). When the relative CD45.1 vs CD45.2 composition for each genotype was displayed on the t-SNE plot, we observed a complete loss of all CD45.2^+^ B-cell progenitor populations in *Mtg16*^*−/−*^ transplanted animals (Fig. 4B, upper right). While the extent to which CD45.2^+^ cells were selected against in *Mtg16*^*F210A*^ mice was less severe when compared with the complete deletion of *Mtg16*, the effect was also observed throughout all Hardy Fractions (Fig. 4B, lower left), suggesting that the impaired ability of *Mtg16*^*−/−*^ and *Mtg16*^*F210A*^ bone marrow cells to give rise to B cells likely preceded B cell lineage commitment. Consistent with results from the spleen, *Mtg16*^*P209T*^ transplanted animals showed levels of CD45.2^+^ B cell populations comparable to wild-type controls (Figure 4B, lower right). The percentage of CD45.2^+^ cells contributing to each Hardy Fraction was quantified for each mouse, and again revealed that in *Mtg16*^*−/−*^ and *Mtg16*^*F210A*^ transplanted animals, there was a reduction in the earliest B cell progenitor population (Hardy Fraction A), that was maintained but not worsened throughout all of B cell development (Fig. 4C, Fig. S1).

**Figure 4.**
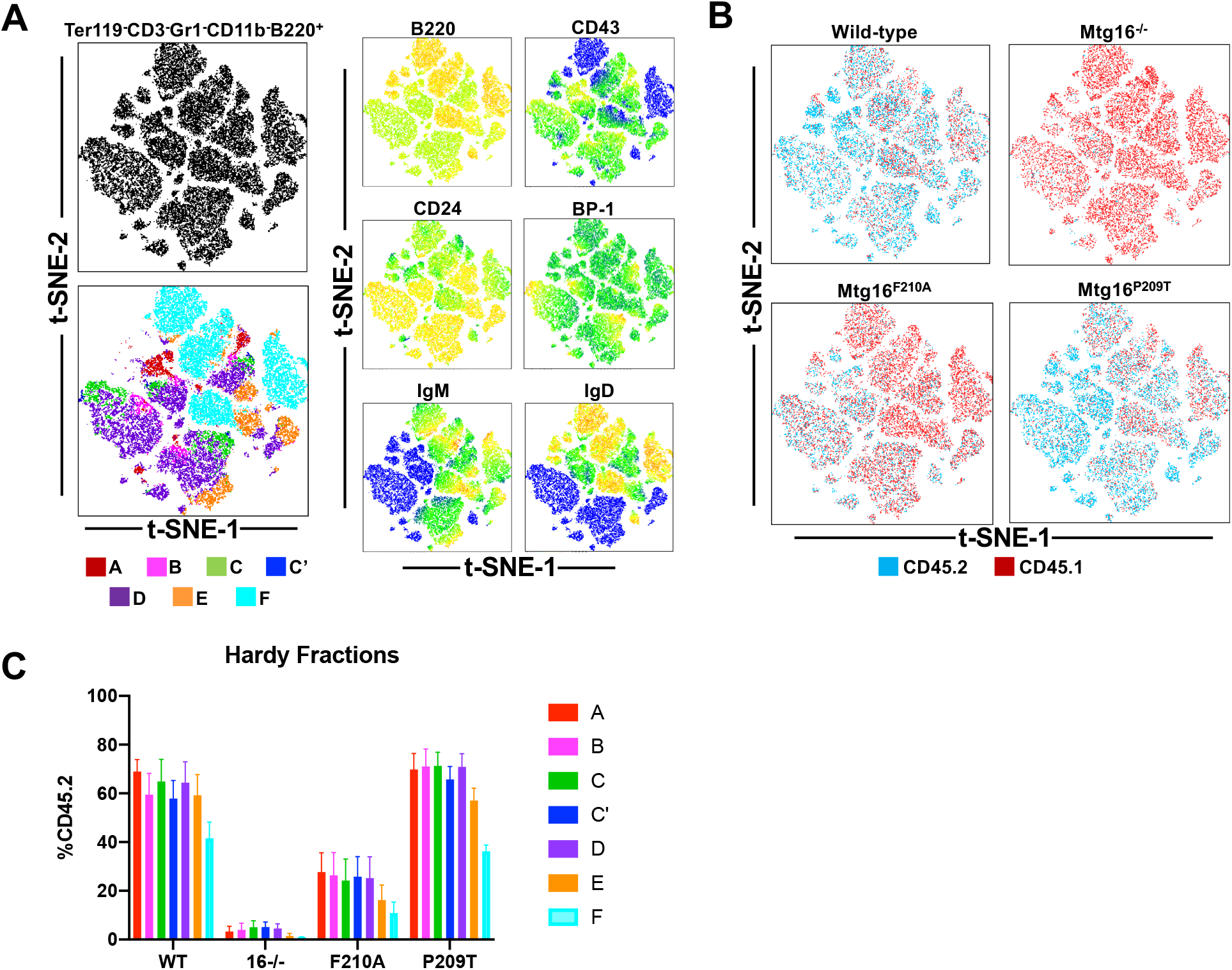
All B cell progenitor populations are reduced in *Mtg16*^*−/−*^ and *Mtg16*^*F210A*^ mice. (A) 3000 lineage-negative, B220+ bone marrow cells from 5 mice of each genotype were concatenated into a single file. T-SNE dimensionality reduction was performed based B cell development parameters and the resultant plot was generated (upper). Manual gating of Hardy Fractions A-F was color coded and displayed on the t-SNE plot (lower). Heatmap depicts the relevant expression of each of these Hardy Fraction surface markers across clusters (right). (B) Genotypes were gated from the concatenated fcs file and the relative contribution of CD45.1+ and CD45.2+ to each cluster was displayed for each genotype. (C) Quantification of gated Hardy Fractions for each genotype is presented (*WT*, n=8; *Mtg16*^−/−^, n=7; *Mtg16F210A*, n=8; *Mtg16P209T*, n=8).

### Mtg16^F210A^ exhibits a reduction in all thymic T cell populations

*Mtg16*^*−/−*^, *Mtg16*^*F210A*^, and *Mtg16*^*P209T*^ transplanted animals all exhibited a reduction in CD3^+^ T cell populations in the spleen. Therefore, we asked whether the loss of Mtg16 function affected T cell development in the thymus. Lineage negative (Gr1^−^, CD11b^−^, Ter119^−^, B220^−^) thymocytes were subject to t-SNE dimensionality reduction based on relevant compensation adjusted parameters (CD3, CD4, CD8, CD147, CD69, TCRβ). Manual gating based on CD4/CD8 staining identified CD4 single positive (SP), CD8 SP, CD4^+^CD8^+^ (double positive, DP), and CD4^−^ CD8^−^ (double negative, DN) populations which were projected onto the t-SNE plot (Figure 5A). Heatmaps depicting additional markers of T cell development and activation were projected onto the t-SNE plot to further refine T cell populations (Fig. 5B).

**Figure 5.**
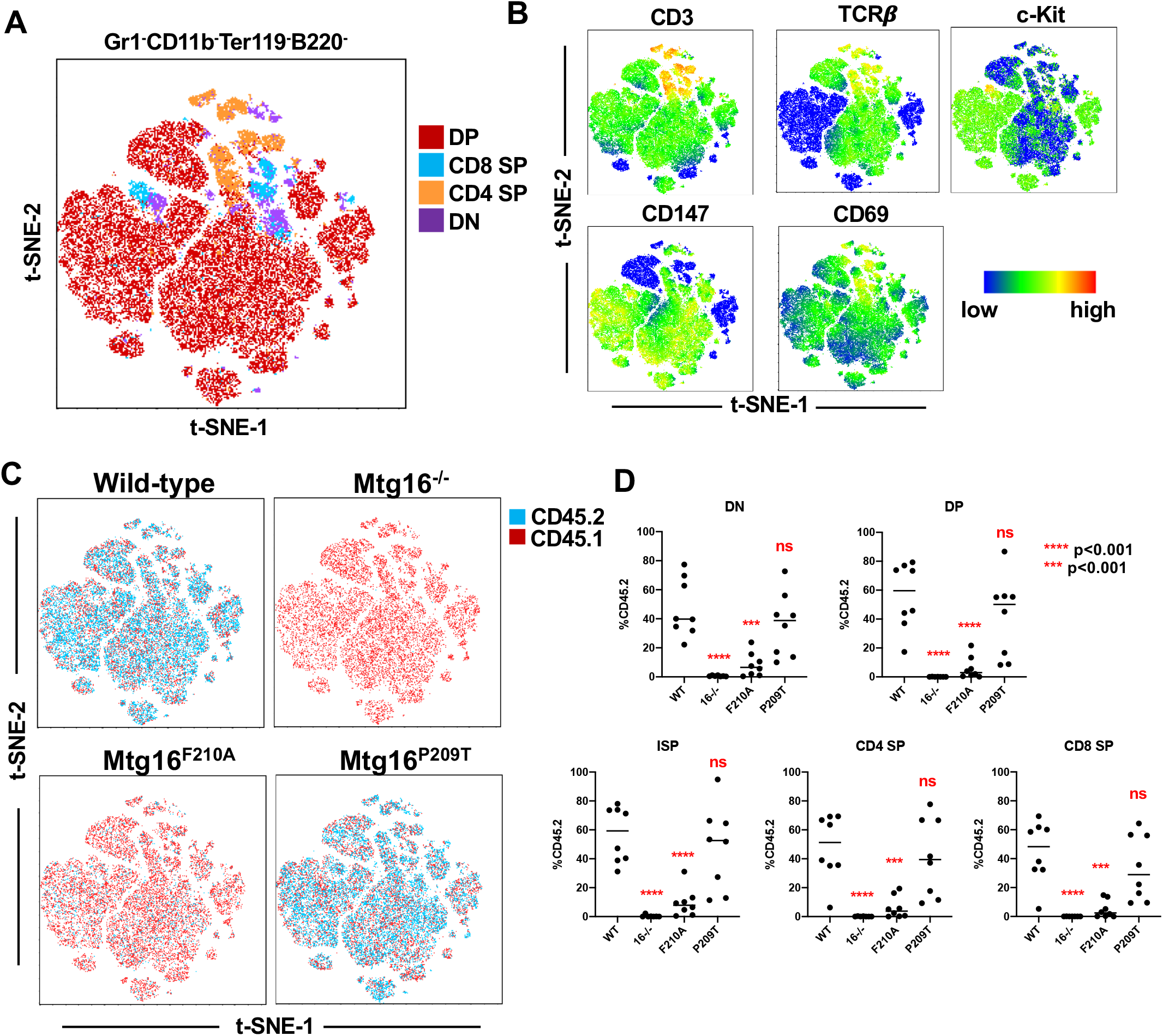
The earliest T cell progenitor populations are reduced in *Mtg16*^*−/−*^ and *Mtg16*^*F210A*^ mice. (A) 3000 lineage-negative (Gr1-CD11b-Ter19-B220-) thymocytes from all mice of each genotype were concatenated into a single file and t-SNE dimensionality reduction performed based on T cell parameters. (B) Heatmaps depict the relative expression of markers associated with T cell development and activation across all clusters. (C) The relative contribution of CD45.1+ and CD45.2+ thymocytes to each cluster is displayed for each genotype. (D) Quantification of each thymocyte population is presented. DN=CD4^−^CD8^−^, DP=CD4^+^CD8^+^, ISP=CD4^−^CD8^+^CD147^+^CD3^−^, CD4 SP=CD4^−^CD8^−^, CD8 SP=CD4^−^CD8^+^CD147^−^CD3^+^.

When relative CD45.1^+^ vs CD45.2^+^ populations were displayed for each genotype, it was clear that *Mtg16*^*−/−*^ marrow was selected against in all thymic populations, regardless of developmental stage (Fig. 5C, upper right). Similar to B cell development in the bone marrow, *Mtg16*^*F210A*^ transplanted animals also showed a reduction in all thymic populations (DN-DP-SP), but to a slightly lesser degree than *Mtg16*^*−/−*^ animals. Surprisingly, even though the CD3+ T cell population was significantly reduced in the spleens of *Mtg16*^*P209T*^ transplanted mice, there was no significant difference in thymic T cell populations (Fig. 5C, lower right, Fig. 5D). The percentage of CD45.2^+^ cells contributing to each thymocyte population was plotted for each mouse. In addition to DN, DP, CD4 SP and CD8 SP, this included immature single positive cells (CD4^−^ CD8^+^CD3^−^CD147^+^), which are cells transitioning from DN to DP (Fig. S2) (Paterson and Williams, 1987). This quantification confirmed a significant reduction of all thymocyte populations in *Mtg16*^*−/−*^ and *Mtg16*^*F210A*^ mice, and no statistically significant decrease in thymocyte populations in *Mtg16*^*P209T*^ mice (Fig. 5D). Of note, we did observe a large variation in the percentage of CD45.2^+^ thymic T cell populations in *Mtg16*^*P209T*^ animals (Fig. 5D), which could suggest an incomplete penetrance of this mutation.

### Mtg16^F210A^ reduced lymphoid progenitor cells in the bone marrow

Given that even the earliest B and T cell progenitor cells were reduced in *Mtg16*^*−/−*^ and *Mtg16*^*F210A*^ transplanted animals, we next sought to characterize earlier stem and progenitor cell populations in the bone marrow to try to pinpoint the exact defect responsible for the reduction in mature lymphocytes. We first quantified mature CD3^+^ T cells, B220^+^ B cells and Gr1^+^CD11b^+^ myeloid cells present in the bone marrow of transplanted mice. Much like mature cell populations observed in the spleen, the percentage of CD45.2^+^ cells was substantially reduced in myeloid, B cell, and T cell subsets in *Mtg16*^*−/−*^ transplanted animals, while *Mtg16*^*F210A*^ transplanted animals showed a significant reduction in lymphoid, but not myeloid cells, and *Mtg16*^*P209T*^ animals only showed a significant reduction in CD3+ populations (Fig. 6A).

**Figure 6.**
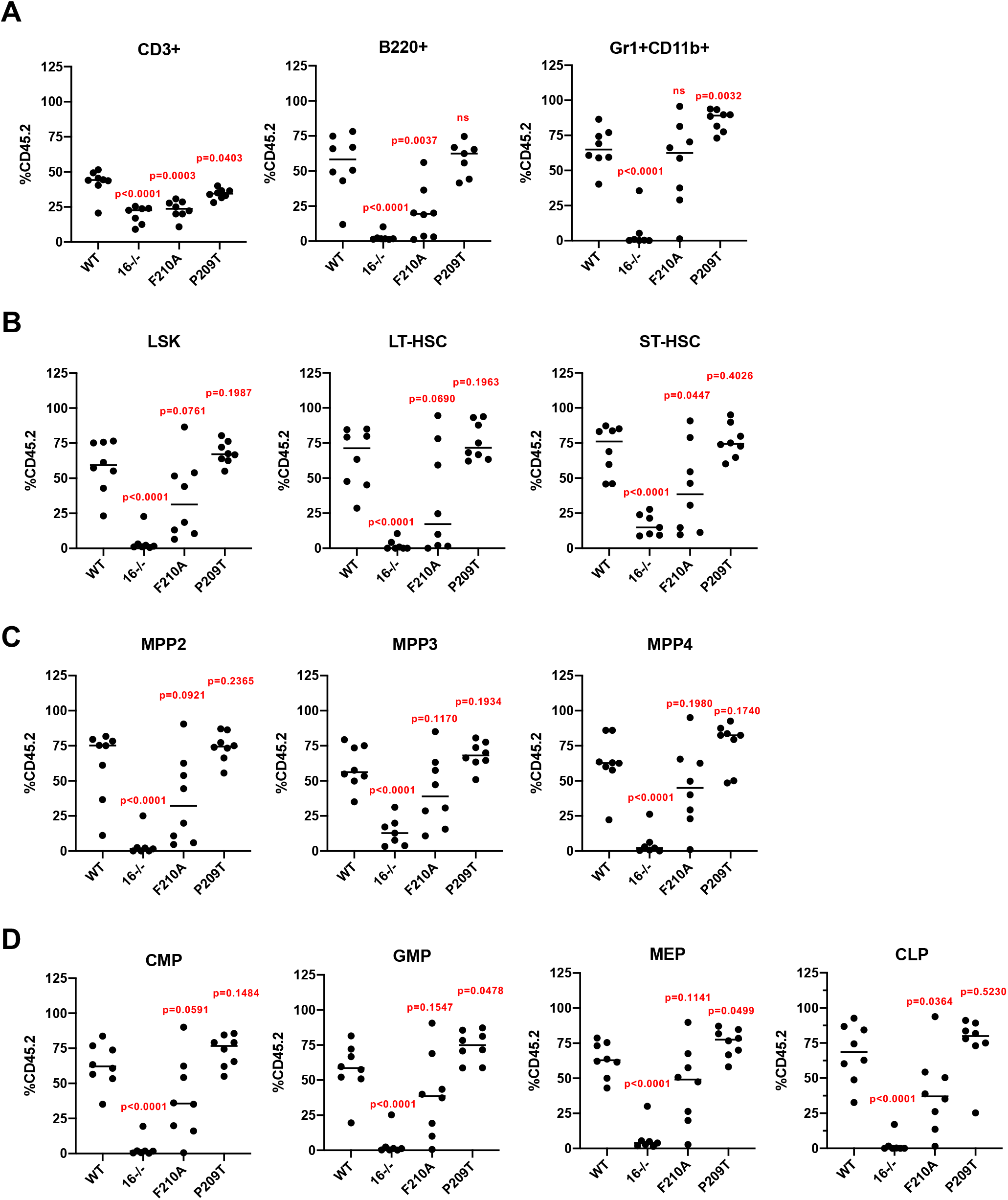
*Mtg16*^*F210A*^ mice show a reduction of common lymphoid progenitors. (A) The percentage of CD45.2^+^ cells contributing to mature T cell (CD3^+^), B cell (B220^+^) and monocyte (Gr1^+^CD11b^+^) populations within the bone marrow were quantified for each genotype. (B-C) The Lineage negative/Sca-1^+^/c-Kit^+^ compartment was further divided into stem and progenitor cell populations on the basis of Flt3 and CD48/CD150 staining. The percentage of CD45.2+ cells contributing to each population was quantified. LT-HSC= Lin−/Sca-1+/c-Kit+/Flt3−/CD150+/CD48-ST-HSC= Lin−/Sca-1+/c-Kit+/Flt3−/CD150−/CD48−. MPP2= Lin−/Sca-1+/c-Kit+/Flt3−/CD150+/CD48+. MPP3= Lin−/Sca-1+/c-Kit+/Flt3−/CD150−/CD48+. MPP4= Lin−/Sca-1+/c-Kit+/Flt3+/CD150−. (D) Myeloid and lymphoid progenitor cell populations were gated and the percentage of CD45.2^+^ cells quantified. GMP= Lin^−^/Sca-1^−^/c-Kit^+^/CD34^+^/CD16/32^hi^. CMP= Lin^−^/Sca-1^−^/c-Kit^+^/CD34^+^/CD16/32^int^. MEP= Lin^−^/Sca-1^−^/c-Kit^+^/CD34^−^/CD16/32^−^. CLP= Lin^−^/Sca^+^1hi/c-Kit^+^/Flt3^+^/IL7R^+^.

We next examined the earliest stem and multipotent progenitor cell populations contained within the Lin^−^cKit^+^Sca-1^+^ (LSK) compartment. Total CD45.2^+^ LSK cells were dramatically reduced in *Mtg16*^*−/−*^ transplanted animals, while a decrease in CD45.2^+^ LSK cells in *Mtg16*^*F210A*^ mice did not reach significance and CD45.2^+^ LSK populations in *Mtg16*^*P209T*^ mice were completely unaffected (Fig. 6B, left). We next employed a gating strategy that more clearly defines functional stem and progenitor cell populations that possess particular lineage biases (Fig. S3) (Pietras et al., 2015). LSK Flt3^−^ cells were further gated based on CD48 and CD150 expression to identify both long-term and short-term stem cell populations (LT- and ST-HSC) as well as MPP2 and MPP3 populations, which are considered myeloid-biased progenitors (Fig. S3). At the same time, we also gated on LSK FLT3^+^CD150^−^ cells, which defines a lymphoid-biased multipotent progenitor population, MPP4 (Pietras et al., 2015). Consistent with previous publications, *Mtg16*^*−/−*^ bone marrow yielded a profound stem cell defect characterized by a dramatic reduction in CD45.2^+^ LT-HSC, ST-HSC, and all MPP populations (Fig. 6B & 6C). Importantly, previous reports have demonstrated that this was not caused by defects in homing or engraftment, but rather a significant loss of the proliferative and self-renewing capacity of *Mtg16*^*−/−*^ stem and progenitor cells (Fischer et al., 2012). In *Mtg16*^*F210A*^ transplants, we observed a large degree of variability for all stem and progenitor populations, which suggested that the *Mtg16*^*F210A*^ mutation was not fully penetrant in earlier stem and progenitor cells. Thus, while we saw a downward trend in CD45.2^+^ LT-HSC, ST-HSC, and all MPP populations in *Mtg16*^*F210A*^ transplanted animals, only the reduction in the ST-HSC population reached significance (Fig. 6B & 6C). Finally, there was no reduction in the CD45.2^+^ population in any stem or multipotent progenitor cell population in *Mtg16*^*P209T*^ transplanted animals (Fig. 6B & 6C), suggesting that an intermediate loss of E protein repression was not sufficient to alter stem or progenitor cell proliferation or self-renewal capacity.

Finally, we examined the effect of Mtg16 mutation on the development of more committed myeloid and lymphoid progenitor cells. Myeloid progenitor populations, defined as Lin^−^cKit^+^ (LK), were further gated based on CD34 and CD16/32 expression (Fig. S3) to identify common myeloid progenitors (CMP), granulocyte/monocyte progenitors (GMP), and megakaryocyte/erythroid progenitors (MEP). We also identified common lymphoid progenitors as Lin^−^/IL7R^+^/FLt3^+^/cKit^+^/Sca^hi^ *Mtg16*^*−/−*^ transplanted animals again exhibited a near complete elimination of CD45.2^+^ cells from all progenitor populations including CMP, GMP, MEP and CLP, while *Mtg16*^*P209T*^ transplanted animals showed no significant reduction of CD45.2^+^ cells in any of these populations (Fig. 5D). *Mtg16*^*F210A*^ transplanted animals again showed high variation in all progenitor cell populations, yet CLPs specifically showed a significant reduction in CD45.2^+^ cells, suggesting that the effect of *Mtg16*^*F210A*^ on lymphoid development becomes clear at the MPP4 to CLP transition, before cells fully commit down the T cell or B cell path.

## Discussion

The MTG family of transcriptional co-repressors are frequently altered in human cancers. In particular, two family members, MTG8/ETO and MTG16/CBFA2T3, are involved in recurrent translocations in myeloid leukemias. The t(8;21) and t(16;21) translocations fuse the DNA binding region of the transcription factor, *RUNX1*, to the majority of *MTG8* or *MTG16,* respectively (Erickson et al., 1992; Gamou et al., 1998; Miyoshi et al., 1993). Furthermore, MTG16 is also targeted by inv(16)(p13.3q24.3) in non-Downs AMKL (Gruber et al., 2012), suggesting that altered MTG function may drive leukemogenesis. However, recent studies have also implicated MTG family members in solid tumors. For instance, *MTG8* was identified as a frequently mutated gene in squamous cell lung carcinoma (Kan et al., 2010) and as a driver mutation in colorectal carcinoma (Sjoblom et al., 2006; Wood et al., 2007). Moreover, the Cancer Genome Atlas (TCGA) has reported SNVs across all three MTG family members in multiple tumor types. The lack of an apparent mutational hotspot suggests that MTG family members may function as tumor suppressors, and this hypothesis is borne out in mouse models in which the loss of either *Mtgr1* or *Mtg16* triggered intestinal tumor formation in both *APC*^*min*^-driven and inflammatory-induced tumor models (Barrett et al., 2011; McDonough et al., 2017; Parang et al., 2016; Parang et al., 2015; Poindexter et al., 2015; Williams et al., 2013).

Here, we sought to determine if SNVs reported across the Mtg16 NHR1 domain are in fact loss-of-function mutations. To this end, we screened these putative NHR1 mutants for the ability to affect E protein binding and Mtg16-mediated transcriptional repression. While most of these SNVs had no effect, we did observe a partial reduction in Mtg16 repressor activity for some variants, the most dramatic of which was *Mtg16*^*P209T*^. However, it is important to note that the effect of the P209T mutation on Mtg16 transcriptional repression was not as severe as total NHR1 deletion or *Mtg16*^*F210A*^, which has been previously shown to disrupt E protein associations and dramatically impair E protein repression. Thus, we believe that most reported NHR1 mutations are merely SNPs and likely do not affect Mtg16 function. The caveat to this conclusion is that we focused our analysis on Mtg16-E protein binding and regulation, and while E proteins appear to be the proteins that most frequently interact with NHR1, we cannot fully rule out that these SNPs disrupt other critical NHR1 contacts to alter specific Mtg16 activities.

While it appeared that most reported variants were not actually functional mutations, it remained possible that the partial repression defect associated with *Mtg16*^*P209T*^ would be sufficient to impact Mtg16-associated phenotypes. Therefore, we utilized CRISPR-based genome editing to create both the P209T and F210A point mutations in mice. Given that *Mtg16*-deletion from bone marrow resulted in profound defects in stem cell quiescence and overall hematopoietic development (Chyla et al., 2008; Fischer et al., 2012), we asked whether these NHR1-associated mutations would result in similar phenotypes. These experiments allowed us to both address whether the *Mtg16*^*P209T*^ mutation is truly loss of function, as well as determine the contribution of Mtg16-mediated repression of E proteins to the overall function of Mtg16 within hematopoiesis.

By and large, the *Mtg16*^*P209T*^ transplanted hematopoietic system resembled wild-type controls, suggesting that this mutation had very little effect on Mtg16 function and biology. The one exception to this was a significant reduction in mature T cells, which indicates that T cells may be sensitive to even modest deregulation of E protein transcriptional networks. Furthermore, the *Mtg16*^*F210A*^ model supports this idea, as complete disruption of E protein repression clearly phenocopied many aspects of the Mtg16 phenotype (though the severity of those phenotypes was reduced), but had the most dramatic effects on lymphoid cell populations (B and T cells).

The idea that altering the ability of Mtg16 to repress E protein-mediated transcription might affect lymphopoiesis is not altogether surprising. E2A is a critical for B cell development (Bain et al., 1994; Zhuang et al., 1994) and has been implicated in lymphoid priming (Dias et al., 2008). In addition, translocations targeting the hematopoietic-specific class II bHLH proteins, TAL1 and LYL1, are associated with T cell malignancies (Begley et al., 1989; Cleary et al., 1988).

Importantly, these proteins require dimerization with E proteins to bind DNA and are found in stable complexes with Mtg proteins. In mouse models, deletion of *Lyl1* resulted in a reduction in the LSK compartment and defect in B cell development, while myeloid lineages remained largely unaffected (Capron et al., 2006). This phenotype appears remarkably similar to the *Mtg16*^*F210A*^ phenotype reported here, wherein we observe a consistent defect in mature lymphoid cell populations and early lymphoid progenitors (CLP), yet effects on myeloid populations including early myeloid progenitors was highly varied. Thus, the ability of Mtg16 to regulate E protein-dependent transcription is critical for normal lymphopoiesis, while NHR1-mediated contacts with other transcription factors may be more important for myelopoiesis.

## Acknowledgements

We thank the members of the Hiebert lab for helpful discussions, reagents and advice. We thank the Vanderbilt Flow Cytometry and Genome Editing Shared Resources. This work was supported by the T. J. Martell Foundation, the Robert J. Kleberg, Jr. and Helen C. Kleberg Foundation, National Institutes of Health grants (RO1-CA178030, RO1-CA164605 and R01-CA64140) and core services performed through Vanderbilt Digestive Disease Research grant (NIDDK P30DK58404) and the Vanderbilt-Ingram Cancer Center support grant (NCI P30CA68485). The project described was also supported by the National Center for Research Resources, Grant UL1 RR024975-01, and is now at the National Center for Advancing Translational Sciences, Grant 2 UL1 TR000445-06. The content is solely the responsibility of the authors and does not necessarily represent the official views of the NIH.

---

**Supplementary Figure 1.**
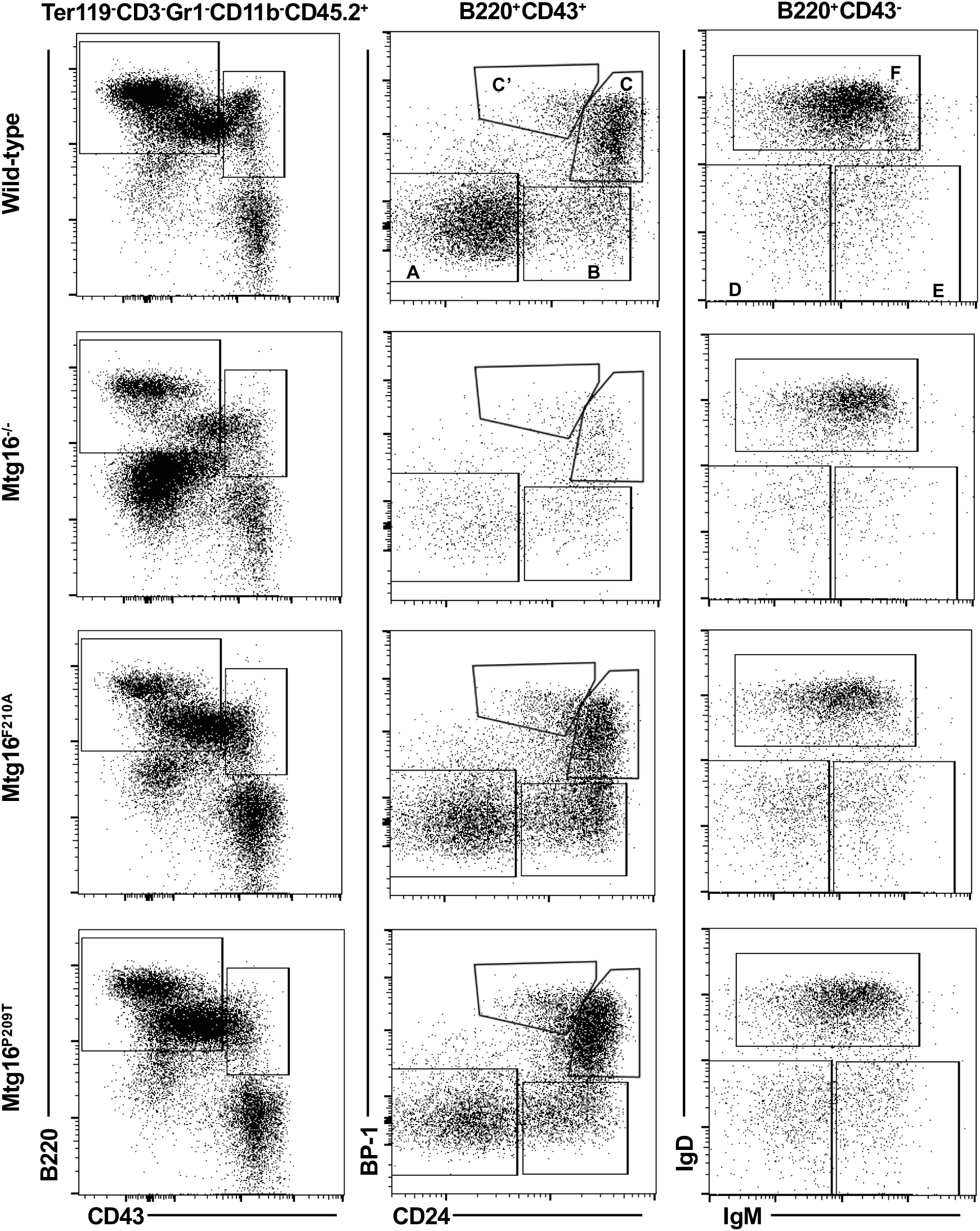
Hardy Fractions monitor B cell development in the bone marrow. CD45.2+Lin− cells from the bone marrow of the indicated mice divided into Hardy Fractions A-F based on the indicated cell surface markers.

**Supplementary Figure 2.**
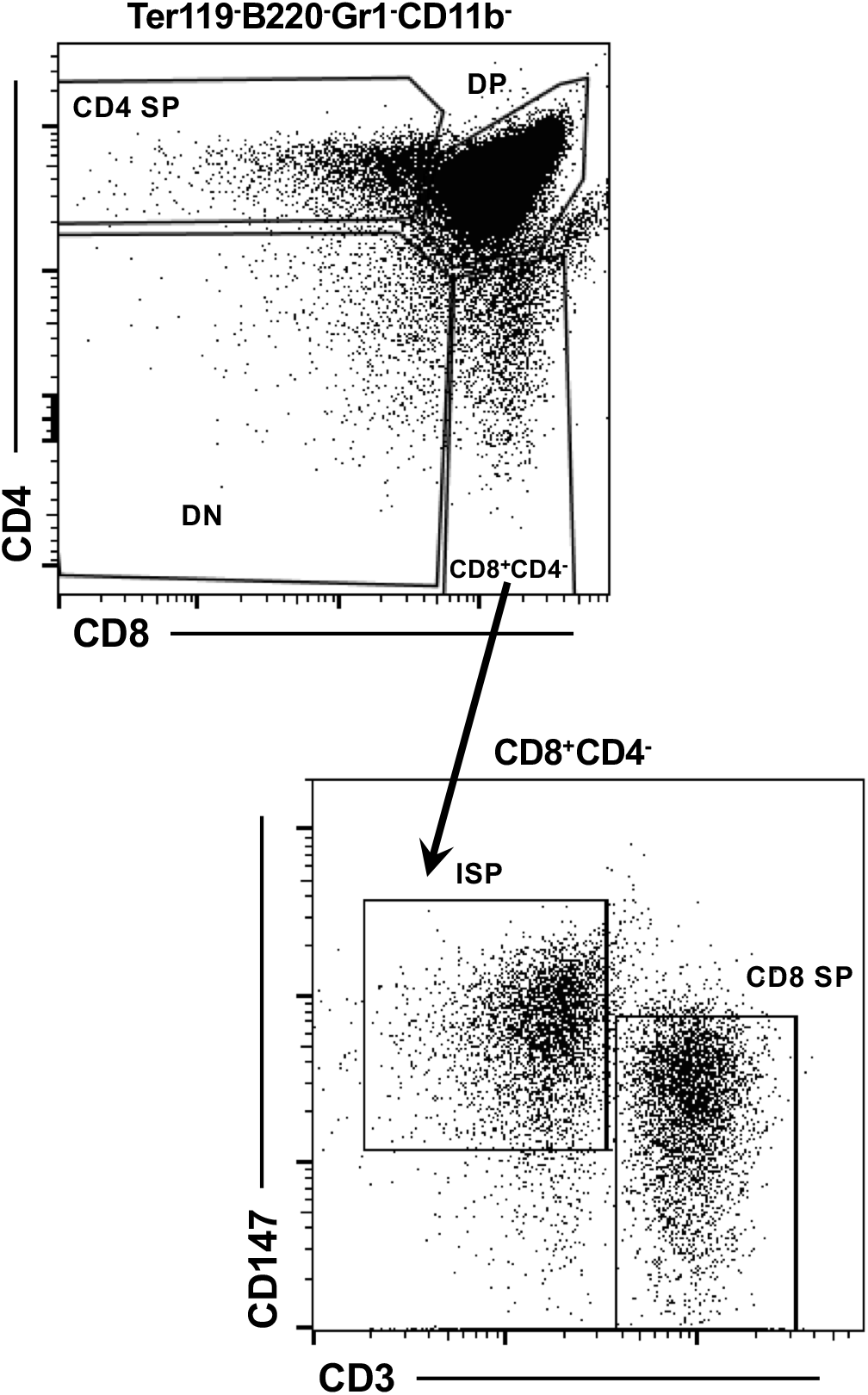
Gating strategy for thymic T cell populations quantified in Figure 5.

**Supplementary Figure 3.**
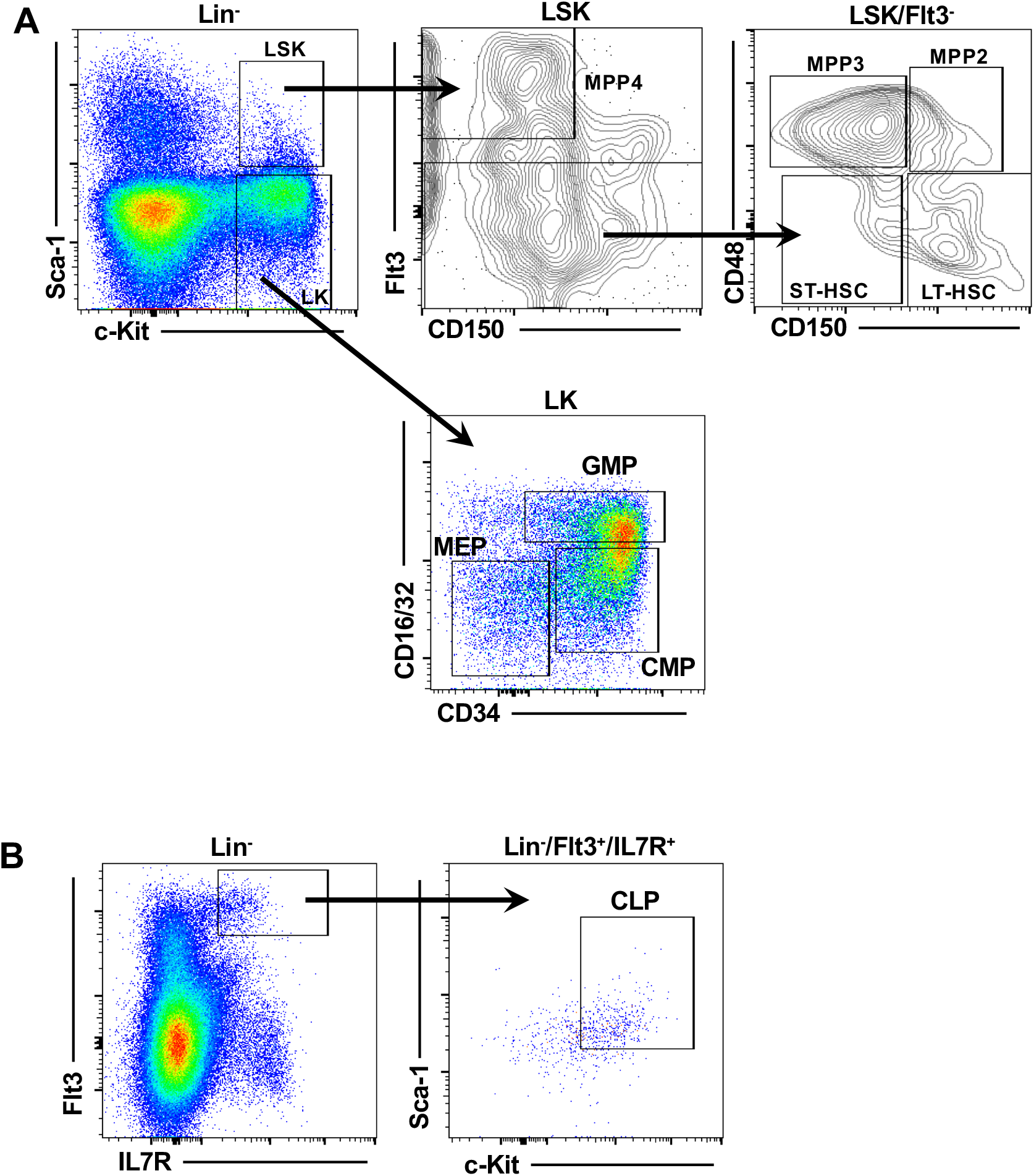
Gating strategy for bone marrow stem and progenitor cell populations which are quantified in Figure 6.

**Supplementary Table 1.**
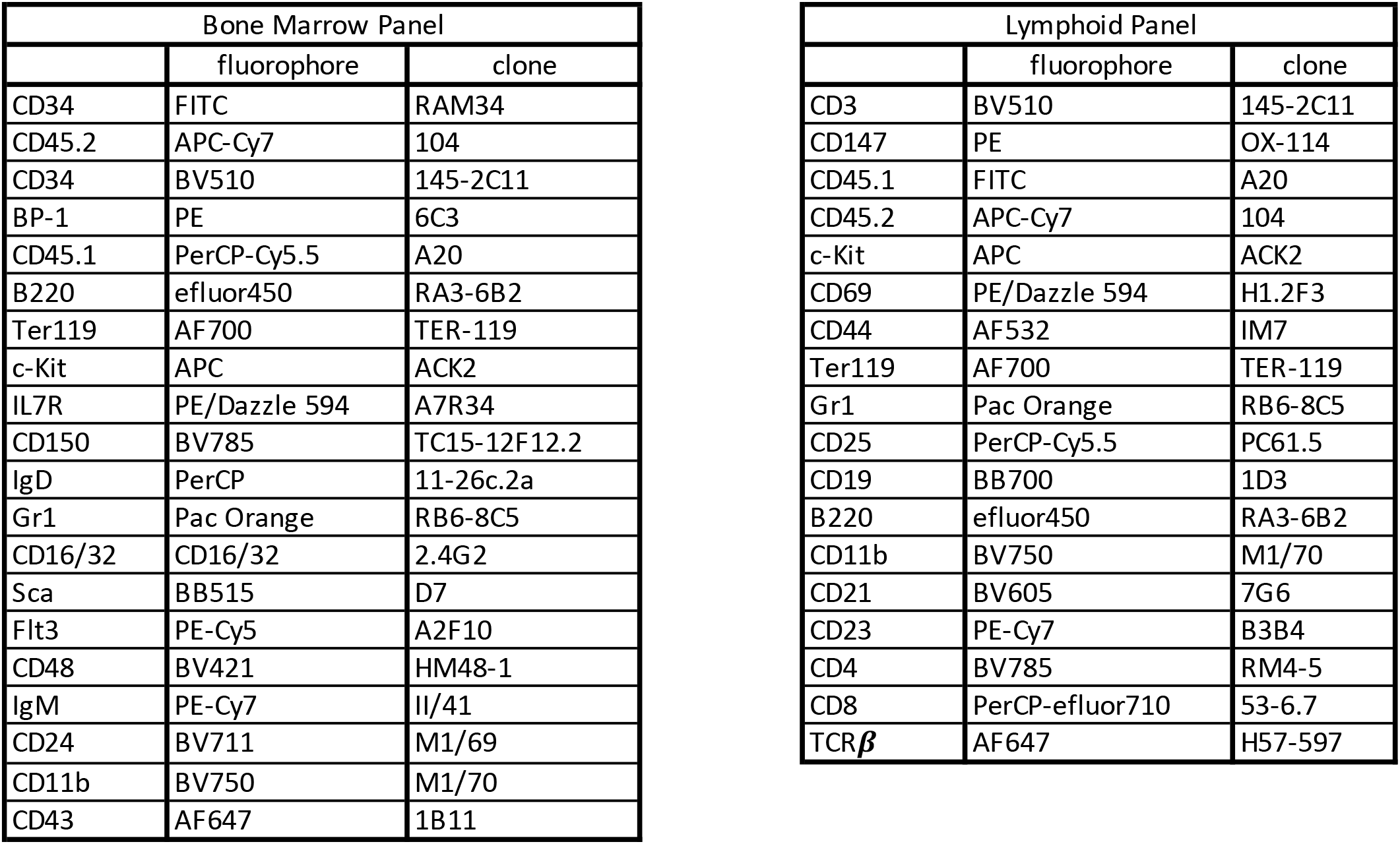
Antibodies used for spectral flow analysis.

| Bone Marrow Panel |  |  |
| --- | --- | --- |
|  | fluorophore | clone |
| CD34 | FITC | RAM34 |
| CD45.2 | APC-Cy7 | 104 |
| CD34 | BV510 | 145-2C11 |
| BP-1 | PE | 6C3 |
| CD45.1 | PerCP-Cy5.5 | A20 |
| B220 | efluor450 | RA3-6B2 |
| Ter119 | AF700 | TER-119 |
| c-Kit | APC | ACK2 |
| IL7R | PE/Dazzle 594 | A7R34 |
| CD150 | BV785 | TC15-12F12.2 |
| IgD | PerCP | 11-26c.2a |
| Gr1 | Pac Orange | RB6-8C5 |
| CD16/32 | CD16/32 | 2.4G2 |
| Sca | BB515 | D7 |
| Flt3 | PE-Cy5 | A2F10 |
| CD48 | BV421 | HM48-1 |
| IgM | PE-Cy7 | II/41 |
| CD24 | BV711 | M1/69 |
| CD11b | BV750 | M1/70 |
| CD43 | AF647 | 1B11 |

**Supplementary Table 1. Antibodies used for spectral flow analysis.**
| Lymphoid Panel |  |  |
| --- | --- | --- |
|  | fluorophore | clone |
| CD3 | BV510 | 145-2C11 |
| CD147 | PE | OX-114 |
| CD45.1 | FITC | A20 |
| CD45.2 | APC-Cy7 | 104 |
| c-Kit | APC | ACK2 |
| CD69 | PE/Dazzle 594 | H1.2F3 |
| CD44 | AF532 | IM7 |
| Ter119 | AF700 | TER-119 |
| Gr1 | Pac Orange | RB6-8C5 |
| CD25 | PerCP-Cy5.5 | PC61.5 |
| CD19 | BB700 | 1D3 |
| B220 | efluor450 | RA3-6B2 |
| CD11b | BV750 | M1/70 |
| CD21 | BV605 | 7G6 |
| CD23 | PE-Cy7 | B3B4 |
| CD4 | BV785 | RM4-5 |
| CD8 | PerCP-efluor710 | 53-6.7 |
| TCRβ | AF647 | H57-597 |

**Supplementary Table 2.**
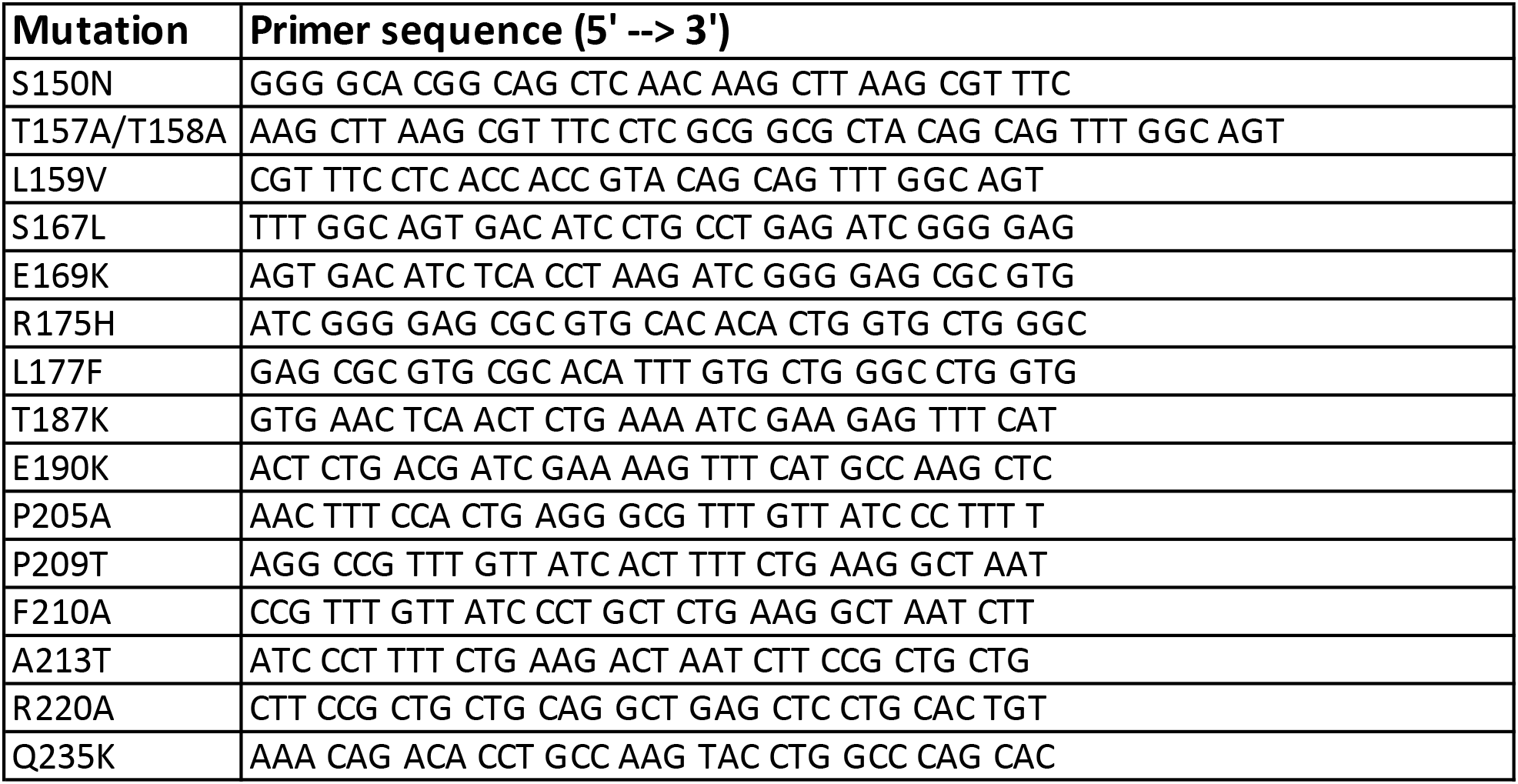
NHR1 mutagenesis primers.

